# Complex Traits and Candidate Genes: Estimation of Genetic Variance Components Across Modes of Inheritance

**DOI:** 10.1101/2022.07.04.498768

**Authors:** Mitchell J. Feldmann, Giovanny Covarrubias-Pazaran, Hans-Peter Piepho

## Abstract

Large-effect loci—those discovered by genome-wide association studies or linkage mapping—associated with key traits segregate amidst a background of minor, often undetectable genetic effects in both wild and domesticated plants and animals. Accurately attributing mean differences and variance explained to the correct components in the linear mixed model (LMM) analysis is important for both selecting superior progeny and parents in plant and animal breeding, but also for gene therapy and medical genetics in humans. Marker-assisted prediction (MAP) and its successor, genomic prediction (GP), have many advantages for selecting superior individuals and understanding disease risk. However, these two approaches are less often integrated to simultaneously study the modes of inheritance of complex traits. This simulation study demonstrates that the average semivariance can be applied to models incorporating Mendelian, oligogenic, and polygenic terms, simultaneously, and yields accurate estimates of the variance explained for all relevant terms. Our previous research focused on large-effect loci and polygenic variance exclusively, and in this work we want to synthesize and expand the average semivariance framework to a multitude of different genetic architectures and the corresponding mixed models. This framework independently accounts for the effects of large-effect loci and the polygenic genetic background and is universally applicable to genetics studies in humans, plants, animals, and microbes.

## Introduction

Today, LMMs are routinely applied in breeding and quantitative genetics research and are used for the prediction of genetic values in plants and animals (VanRaden 2008; Hayes *et al*. 2009; Albrecht *et al*. 2011; Endelman 2011; Crossa *et al*. 2014; Meuwissen *et al*. 2016), or polygenic risk scores (PRSs) in humans (de los Campos *et al*. 2010; Dudbridge 2013; Wray *et al*. 2019; Truong *et al*. 2020; de Los Campos *et al*. 2013; Lello *et al*. 2018, 2019), to estimate the heritability of traits in target populations (Visscher *et al*. 2006, 2008; de los Campos *et al*. 2015; Lehermeier *et al*. 2017; Legarra 2016), and to estimate ecological and evolutionary genetic parameters of behavioral traits (Walsh and Lynch 2018; Walsh *et al*. 2020; Oldroyd 2012; Hemani *et al*. 2013; Ariyomo *et al*. 2013). Genetic values are constructed from a combination of genetic effects; including Mendelian factors; which may have both additive effect and dominance deviations (Pincot *et al*. 2018, 2022), oligogenic factors consisting of few genetic factors and their epistatic interactions appropriate for marker-assisted prediction (MAP) (Tang *et al*. 2006), a polygenic term consisting of a dense genome-wide framework of markers assumed to have minor effects appropriate for genomic prediction (GP); which may also account of additive and dominance sources of variance (Pincot *et al*. 2020; Brandariz and Bernardo 2019), and a residual genetic term consisting of all genetic effects not accounted for by the previous genetic factors (Rutkoski *et al*. 2014; Rice and Lipka 2019; DeWitt *et al*. 2021). The ultimate objective in breeding applications is, typically, predicting the genotypic value, e.g., breeding value or genetic merit of a candidate individual (Knapp 1998; Piepho *et al*. 2008; Piepho 2009; VanRaden 2008; Luby and Shaw 2001; Collard and Mackill 2007). For loci to provide actionable gains or diagnoses, they must explain a significant proportion of phenotypic and genetic variation in a population with alleles in segregation at target loci.

Candidate gene discovery through genome-wide association studies (GWAS) and quantitative trait locus (QTL) mapping is prolific in plant and animal populations (Lander and Botstein 1989; Lander and Schork 1994; Visscher *et al*. 2012, 2017; Korte and Farlow 2013; Yu *et al*. 2006). Despite decades of directional selection in many plant populations, loci impacting traits of interest still segregate, even in advanced breeding materials, and these genome-wide analyses have succeeded in implicating numerous genes and genomic regions in the control of a wide variety of both simple and complex traits (Tang *et al*. 2006; Pincot *et al*. 2018; Wassom *et al*. 2008; Demmings *et al*. 2019a; Rutkoski *et al*. 2014; Rice and Lipka 2019; DeWitt *et al*. 2021; Han *et al*. 2018; Xin *et al*. 2020; Kim and Reinke 2019; Gage *et al*. 2020; Visscher *et al*. 2012, 2017; Andersson 2001; Hayes and Goddard 2001; Anderson *et al*. 2007; Septiningsih *et al*. 2009; Hayes *et al*. 2010; Saatchi *et al*. 2014; Seabury *et al*. 2017), although the utility of such marker-trait associations may not be fully realized (Bernardo 2004, 2016). Large-effect and statistically significant loci typically only explain a fraction of the genetic and phenotypic variance in a population (Feldmann *et al*. 2021), along with the polygenic fraction (Feldmann *et al*. 2022), except in extreme scenarios when Mendelian factors wholly control a trait.

Discovered loci rarely, if ever, explain 100% of the genetic variance, and understanding the multiple sources of variation and how they relate can help breeders and research prioritize targets and mitigate risk (Bernardo 2004, 2014). Genes with significant effects often dominate the ‘non-missing heritability,’ but they can also mask or obscure the effects of other quantitatively acting genes and pleiotropically affect multiple quantitative phenotypes (Mackay 2001; Mackay *et al*. 2009; Lorenz and Cohen 2012; De Villemereuil *et al*. 2018; Eichler *et al*. 2010). For example, mutations in the *BRCA2* gene can have large effects, but be incompletely penetrant, interact with other genes, and may be necessary but insufficient for predicting breast, ovarian, and other cancer risks in women (Gaudet *et al*. 2010). Accurately partitioning the Mendelian, oligogenic, and polygenic sources of variance allows researchers to assess how much value, or risk, specific loci confer.

Here, we use simulations to show that the ASV provides accurate variance component estimates (VCEs) and variance component ratios for all relevant genetic terms regardless study design or population type, e.g., outbred or inbred. We sought to marry the our previously published works (Piepho 2019; Feldmann *et al*. 2021, 2022) and to present a fully realized ASV approach for typical LMM analyses in human, plant, animal, and microbial genetics. We demonstrate how these models can be extended to handle more complex genetic structures, including adding multiple explanatory loci and marker-marker interactions, incorporating non-additive dominance and epistasis variance, and partitioning marker variance into additive and dominance components. We provide examples of expressing the different models and extensions in the freely available `sommer` R package (Covarrubias-Pazaran 2016). We believe that the average semivariance is a powerful tool for answering these questions regardless of the organism, population, or trait.

### Linear mixed model analysis and the average semivariance

The average semivariance (ASV) estimator of total variance (Piepho 2019) and the variance of single markers and marker-marker interactions (Feldmann *et al*. 2021) is half the average total pairwise variance of a difference between entries and can be decomposed into independent sources of variance, e.g., genetic and residual. In this article, we assume that researchers are able to independently replicate entries—as in clonally propagated or inbred crop species—or can collect repeated measures on entries (e.g., individuals, families, or strains)—as in humans and animals—and then estimate the least square means (LSMs), best linear unbiased estimators (BLUEs), or other adjusted entry means in the first stage of a two-stage analysis (Piepho *et al*. 2012; Schulz-Streeck *et al*. 2013; Damesa *et al*. 2017, 2019).

The key idea here is that the adjust entry means, in general, are considered the “phenotype” since we assume independent replication. In animal breeding, “de-regressed” best linear unbiased predictors (BLUPs) are used in GBLUP and GWAS analysis (Strandén and Mäntysaari 2010; Ricard *et al*. 2013; Calus *et al*. 2016; Konstantinov and Goddard 2020). The two-stage approach is commonly applied for GWAS and GP studies in plants (Pincot *et al*. 2018, 2020; Damesa *et al*. 2017; Dias *et al*. 2020; Gogel *et al*. 2018). For simplicity in our demonstration, we assume that the error variance of the observation is 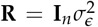, where *n* is the number of entries (e.g., individuals, accessions, genotypes, lines, or animals). The more general approach is to assume a general variance-covariance matrix **R** and, importantly, the average semivariance can efficiently deal with more general forms of **R** and integrated directly into single-stage or multi-stage analyses. We explore ASV in a fully efficient two-stage analysis below in this article.

The form of the linear mixed model (LMM) for this analysis assuming only one explanatory marker is:

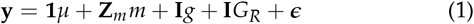

where *y* is the vector of LSMs with *y* ~𝒩 (*µ*, **V**), *µ* is the population mean and the only fixed effect, *m* is the random effect of the main-effect locus with 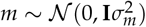, *g* is the random additive genetic effect associated with the genome-wide framework of marker excluding *m* with 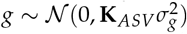, *G*_*R*_ is the random residual genetic term—the portion of the total genetic effect not accounted for by *m* or *g*—with 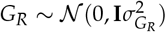, and *ϵ* is the random residual term with *ϵ* ~ 𝒩 (0, **R**). We then calculated **K**_*ASV*_ as:

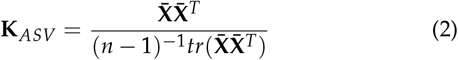

where 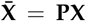 is the mean-centered marker matrix, 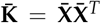 is the realized genomic relationship or kinship matrix, 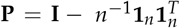is the idempotent mean-centering matrix, and *tr*(.) is the trace. **Z**_*m*_ is a *n×n*_*m*_ dimension design matrix linking levels of the explanatory locus to LSMs in *y*, where *n*_*m*_ is the number of marker genotypes.

The ASV definition of total variance from LMM (1) is:

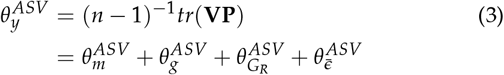

where 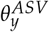 is the total phenotypic variance, **V** is the variancecovariance among observations, 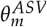 is the average semivariance of the simple genetic term, 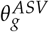 is the average semivariance of the polygenic term, 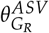 is the average semivariance of the residual genetic term, and 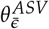 is the average semivariance of the residuals.

The ASV definition of the genomic variance is:

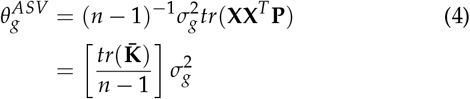

In general, we replace the unknown parameter values 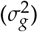 with their REML estimates 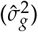 to obtain the ASV estimates 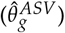. Following this form, it is possible to extend LMM (1) to include dominance and epistatic sources of variance (see below). The ASV definition of the marker associated genetic variance is:

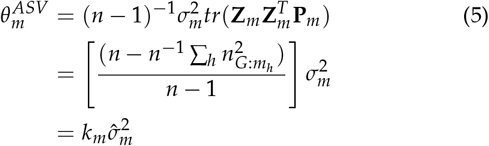

It is possible to extend this using the approach for multi-locus models as in (8), with and without marker-marker interactions, described in (Feldmann *et al*. 2021). The ASV definition of the residual genetic variance is:

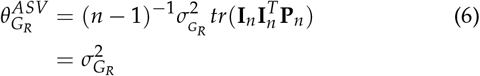

Importantly, all terms are estimated on the same scale as the residual variance 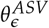and are estimates on an entry-mean basis. The ASV definition of the residual variance is:

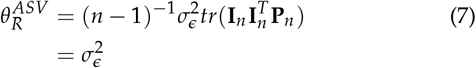

### Linear mixed model extensions incorporating the average semivariance

While an important model, LMM (1) only covers a narrow scope of the possible genetic models and experiments that might exist, and we want to provide researchers with a clear strategy for expanding this approach to more complex systems. This section demonstrates how to partition the additive and dominance variance from a single marker, incorporate multiple explanatory loci, their interactions into the model, and non-additive polygenic terms, and achieve a fully efficient two-stage analysis. Depending on the population, trait, environment, etc. the unique components of the models demonstrated here can be hybridized and merged to accurately and holistically decompose the multitude of potential sources of genetic variation. The code to execute these models using the sommer v4.1.7 (Covarrubias-Pazaran 2016) is provided in the methods.

#### Extension #1: Incorporating multiple target loci and locus-locus interactions

It is common for multiple QTL to be implicated from genetic stud-ies (Tang *et al*. 2006; Rutkoski *et al*. 2014; Vasconcellos *et al*. 2017; Lopdell *et al*. 2019; Legare *et al*. 2000; Cockerton *et al*. 2019; Rice and Lipka 2019; Demmings *et al*. 2019b), the utility of which is not always certain (Bernardo 2001, 2004). While the simulations in this paper rely exclusively on LMM (1), this model can be easily expanded to include multiple explanatory loci and their interactions or statistical epistasis (Moore and Williams 2005; Álvarez-Castro and Carlborg 2007), as demonstrated by (Feldmann *et al*. 2021). For example, the LMM with three main-effect loci, denoted *m*_1_, *m*_2_, and *m*_3_, is:

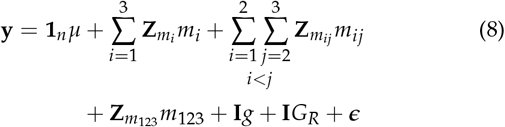

where *m*_*i*_ is the random effect of the *i*-th main-effect marker, *m*_*ij*_ is the random effect of the two-way interaction between the *i*-th and *j*-th markers, and *m*_123_ is the random effect of the three-way interaction between the three main-effect loci. 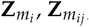, and 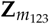 are design matrices that link levels of the explanatory marker and interactions to LSMs in *y*. The rest of the terms have the same definitions.

#### Extension #2: Partitioning 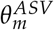 into additive 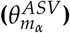 and dominance 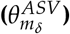 components

The factor coding of the Mendelian and oligogenic markers is a different approach than is standard in GWAS (Korte and Farlow 2013; Visscher *et al*. 2012, 2017). In GWAS, markers are typically treated as fixed and coded numerically, e.g., the dosage model. Assuming that a researcher is working with an outbred species (*H* ≠ 0), the dominance deviations can be significant, and partitioning the additive and dominance sources of variance from significant markers can be helpful in hybrid crop breeding and disease risk prognoses. Our goal is to partition 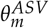into its additive 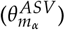 and dominance 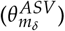 components.

Here, we demonstrate an LMM that can be used to partition the additive and dominance sources of variance of the main effect marker. The form of the linear mixed model (LMM) for this analysis assuming only one explanatory marker is:

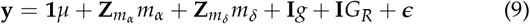

where *m*_*α*_ is the random effect of the main-effect locus with 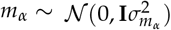 and *m*_*δ*_ is the random effect of the main-effect locus with 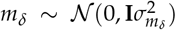. 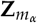is an *n* × 3 design matrix linking marker genotypes to observations and 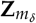 is an *n ×* 2 design matrix linking genotypic state, either homozygous (*AA* and *aa*) or heterozygous (*Aa*), to observations. Other terms are as defined in LMM (1).

The average semivariance associated with *m*_*α*_ is obtained as in (5) by:

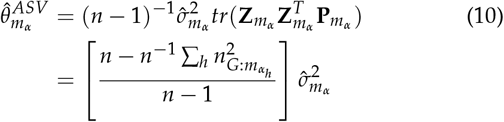

where 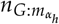 is the number of entries nested in the *h*-th marker genotype (Feldmann *et al*. 2021). The average semivariance associated with *m*_*δ*_ is obtained by :

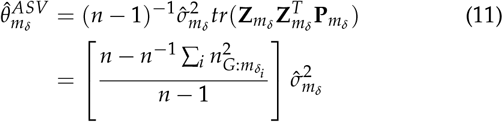

where 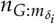 is the number of entries nested in the *i*-th genetic state. The sum of 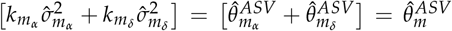 and 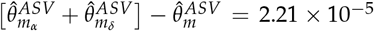. 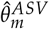 is an unbiased estimate of the variance explained by a marker (Feldmann *et al*. 2021). The likelihood ratio (LR) between LMM (1) and (9) was *LR* ≈ 0 and was not significant in any simulated populations (*P*_*LR*_ > 0.2), suggesting that there is no appreciable difference between the model likelihood of (1) and (9). The same marker variance is estimated in both LMMs, (1) and (9), and the estimates are equal. Note that we were not able to fit LMM (9) in all software and had to use either `sommer::mmer()` or `asreml::asreml()`.

#### Extension #3: Incorporating additional polygenic terms for dominance (*g*_*δ*_) deviations

LMM (1) can also be extended to include both additive (*g*_*α*_) and dominance (*g*_*δ*_) sources of genomic variance (Vitezica *et al*. 2013; Kumar *et al*. 2015; Vitezica *et al*. 2017; Xiang *et al*. 2018; Sun *et al*.2014; Ali *et al*. 2020; Zhang *et al*. 2021; Martini *et al*. 2016). The form of the LMM for analysis with both *g*_*α*_ and *g*_*δ*_ assuming only one explanatory marker *M* is:

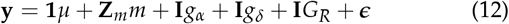

where *g*_*α*_ and *g*_*δ*_ are random effect vectors for the additive and dominance polygenic effects, respectively, with 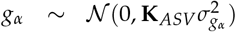 and 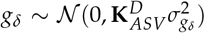. The average semivari-ance dominance kernel is:

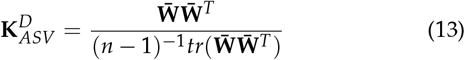

where **W** = 1 − |**X**|, assuming **X** is coded [-1,0,1], and 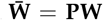. This is a feasible approach to improve genetic performance in crossbred populations with large dominance genetic variation (Nishio and Satoh 2014; Vitezica *et al*. 2017; Xiang *et al*. 2018; Wolfe *et al*. 2021). Both **K**_*ASV*_ and 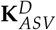have the matrix properties proposed by Speed and Balding (2015); i.e., *n*^−1^*tr*(**K**) = 1 and *n*^−2^ Σ_*i*_ Σ_*j*_ *K*_*ij*_ = 0. Not surprisingly, the dominance variance estimated with 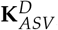were accurate and the relative bias from 100 simulated populations was −3.32%.

Further extensions for additive-by-additive *A* × *A* or additive-by-dominance *A* × *D* polygenic interactions are also possible (Nishio and Satoh 2014; Covarrubias-Pazaran 2016; Vitezica *et al*. 2017). These matrices are often calculated as the Hadamard product (element-wise multiplication, °) of **K**_*ASV*_ and/or 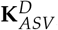, where the additive-by-additive epistasis GRM is 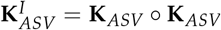. This matrix has the same essential properties as **K**_*ASV*_, and so we hypothesize that the ASV estimted variance components will be accurate for these terms as well.

#### Extension #4: Stage-wise LMM analysis for multi-environment trials (METs) and meta-analysis in plant breeding

Two-stage, or stage-wise, analyses are the *status quo* in plant breeding trials in both academic studies and seed industry (Piepho *et al*. 2012; Damesa *et al*. 2017, 2019; Endelman 2022). The reason for this is that plant breeders are often not interested in the performance *per se* of a line or hybrid *within* a specific location, unless the presence of cross-over (rank change) *G* × *E* is very large enough to make data from one target environment non-informative in another target environment. Instead, plant breeders are often more interested in the ranking and performance of entries averaged across all environments (Bernardo 2020). It is common then to fit a first model that accounts for the variation of random design elements, e.g., locations, years, blocks, and fixed genotype effects to obtain the estimated marginal means (EMMs) or best linear unbiased estimators (BLUEs) as adjusted entry means. These adjusted entry means are then used as the phenotype or response variable in GWAS and genomic prediction studies. However, the naive approach is not “fully efficient” (Piepho *et al*. 2012) and assumes that adjusted entry means are IID; i.e., 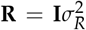. However, due to incomplete block and augmented designs, missing data, and changes in experiment designs over time and location, IID entry means are rarely observed in practice. To fully utilize the data, however, the variance-covariance matrix of the estimates from Stage 1 must be included in Stage 2 (Piepho *et al*. 2012; Damesa *et al*. 2017), which is not possible with many software packages for genomics-assisted breeding.

The LMM for stage one is:

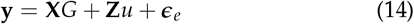

where **X** is the fixed effect design matrix linking observations to entries, **Z** is the random effect design matrix for design (e.g., blocks) elements within each environment (e.g., years and locations), and *ϵ*_*e*_ are the residuals and *ϵ*_*e*_~ 𝒩 (0, **R**_*e*_), where **R**_*e*_ is the residual variance-covariance matrix estimated in the *e*-th environment. **R**_*e*_ can be estimated with or without spatial or autoregressive correlations (Farfan *et al*. 2015; Rodríguez-Álvarez *et al*. 2018; Anderson *et al*. 2018; Selle *et al*. 2020). This model is fitted for each environment independently. From these models, we obtain the adjusted entry means 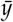 and the residual variance covariance matrices **R**_*e*_ from each of *e* = 1,…, *n*_*e*_ environments, where *n*_*e*_ are the number of environments. For CRD or experiments without design elements the obtained variance-covariance matrix will be diagonal. Assume that we have two environments, we will obtain **R**_1_ from environment 1 and **R**_2_ from environment 2. We can then construct the 2*n* × 2*n* block-diagonal stage-one **Ω** matrix as:

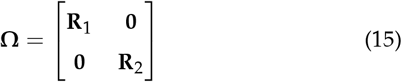

This block-diagonal form indicates that the residuals among entries are uncorrelated among environments. For simplification, **R**_*e*_ can be approximated by diagonal matrices in several different ways (Smith *et al*. 2001; Möhring and Piepho 2009; Welham *et al*. 2010; Piepho *et al*. 2012; Moehring *et al*. 2014), but here we use to the full variance-covariance matrix from each experiment *e*. Importantly, we need to carry **Ω** over from the stage-one analyses to stage-two of the analysis.

The LMM for stage two is then:

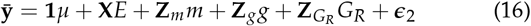

where 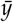 are the adjusted entry means from stage-one, *µ* is the population mean, **X** is the fixed effect design matrix linking environments to adjusted entry means, *E* are the fixed environmental effects, *g* is the random additive genetic effect associated with the genome-wide framework of marker excluding *m* with 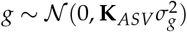, *G*_*R*_ is the random residual genetic term—the portion of the total genetic effect not accounted for by *m* or *g*— with 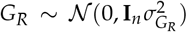, and *ϵ*_2_ is the structured residual term from stage-one with *ϵ*_2_ ~ 𝒩 (0, **Ω**) This approach is accessible to researchers via the sommer, asreml, and StageWise packages in R (Covarrubias-Pazaran 2016; Butler 2021; Endelman 2022) and in SAS.

We created 100 simulated population (*n* = 1, 000; *m* = 5, 000) using a similar approach to the other simulations in this experiment. However, in this experiment we included Environmental and Block within Environment effects. We estimates the variance explained by the polygenic background, a large effect locus, the residual genetic variance, and non-genetic residual. The single stage analysis yielded relative biases of *−* 0.67%, *−* 0.33%, *−* 0.67%, and 0.41% for the marker variance 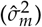, genomic variance 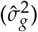, residual genetic variance 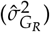, and residual variance 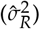, respectively (Fig 3). The two stage analysis yielded relative biases of *−*0.83%, *−*4.08%, 0.15%, and 0.16% for the marker variance 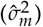,genomic variance 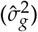, residual genetic variance 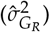, and resid-ual variance 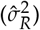, respectively (Fig 3).

#### Extension #5: Incorporating *k*_*M*_ directly into LMM analyses

In (Feldmann *et al*. 2021), we introduced ASV into LMMs for individual markers in genetic analysis as a *post hoc* adjustment of the variance explained by a marker by *k*_*M*_(5). This directly led to (Feldmann *et al*. 2022), in which we showed that ASV estimates of the genomic variance could be obtained by scaling the genomic relationship prior to the LMM analysis and introduced **K**_*ASV*_, eliminating the need for any *post hoc* adjustment. Using statistical packages such as sommer (Covarrubias-Pazaran 2016), we can directly apply *k*_*M*_ to the variance-covariance matrix for large effect loci *M* and their interaction in our model. Typically, the identity matrix is used as the variance-covariance matrix and levels of the random effect are assumed to have the same variance and no covariance. In (Feldmann *et al*. 2021) we multiplied the average marginal variance component by *k*_*M*_ to obtain the ASV component. Instead, if we define 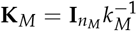, where **K**_*M*_ is *n*_*M×*_*n*_*M*_ and *n*_*M*_ is the number of marker genotypes. We can essentially think of **K**_*M*_ in the same way that we think of genomic relationship matrices; e.g., **K**_*ASV*_, except that we apply **K**_*M*_ to the levels of the marker genotype instead of entries. With this approach, we maintain the levels of the factor come from the same variance and zero covariance, but our scaling factor embedded directly in the model eliminating the need for adjustment. Embedding *k*_*M*_ in the LMM analysis using **K**_*M*_ is equivalent to the *post hoc* adjustment that we proposed in (Feldmann *et al*. 2021), and so it is up to the user to determine which approach they prefer.

## Results and Discussion

### Candidate Genes and Complex Traits

Bernardo (2014) was the first to propose an integration of MAP and GP and since then empirical studies have validated the methodology (Rutkoski *et al*. 2014; Zhang *et al*. 2014; Rice and Lipka 2019; Spindel *et al*. 2016) while others have shown little-to-no improvement over GP (Li *et al*. 2015; Galli *et al*. 2020), suggesting that modeling significant markers can improve prediction accuracy only when markers explain a *significant* portion of both genetic and phenotypic variance (Galli *et al*. 2020). With the high densities of genome-wide markers commonly assayed in gene finding studies, investigators often identify markers tightly linked to candidate or known causal genes as exemplified by diverse real world examples (Andersson 2001; Hayes and Goddard 2001; Anderson *et al*. 2007; Gaudet *et al*. 2010; Hayes *et al*. 2010; Jensen *et al*. 2012; Visscher *et al*. 2012; Septiningsih *et al*. 2009; Saatchi *et al*. 2014; Visscher *et al*. 2017; Freebern *et al*. 2020; Li *et al*. 2021; Korte and Farlow 2013). The candidate marker loci are nearly always initially identified by genome-wide searches using sequential (marker-by-marker) approaches such as GWAS and QTL analysis. Following the discovery of statistically significant marker-trait associations from a marker-by-marker genome-wide scan, the natural progression would be to analyze single- or multi-locus genetic models where the effects of the discovered loci are simultaneously corrected for the effects of other discovered loci, e.g., polygenic variation (Stroup *et al*. 2018; Gbur *et al*. 2020).

A marker will not explain a large portion of variance if that marker does not have a large, detectable effect and, thus, markers that explain a large portion of genetic variance will be the most useful for MAP. For example, consider Fusarium wilt resistance in strawberry which is conferred by a single dominant acting locus *Fw*1 (Pincot *et al*. 2018, 2022). This locus explains nearly 100% of both the phenotypic and genetic variance and the mean differences delineate resistant vs susceptible genotypes, and thus there is almost no added benefit of a genome-wide sample of markers over the single-marker assay (*m*) for product delivery and germplasm improvement. While variance explained is directly linked to the effect size, it is not a direct substitute. However, the random effect machinery allows for researchers to obtain variance component estimates and effect sizes (e.g., BLUPs) simultaneously (Searle *et al*. 1992) eliminating the need for multiple statistical models to assess the variance explained and the effect size of a target locus. The BLUP procedure is directly applied in this model, so it is natural to use the same statistical machinery to estimate GEBVs by GBLUP and the genetic effect of a locus.

As a point of contrast, yield in maize (*Zea mays*) is heritable but no single locus explains any appreciable amount of phenotypic or genotypic variance (Heffner *et al*. 2009, 2010; Yang *et al*. 2017; Brandariz and Bernardo 2019; Gage *et al*. 2020; Zhang *et al*. 2019). For improvement of yield in maize, GP is potentially a more valuable approach because the researcher, or breeder, can predict the polygenic value (*g*) without relying on any one particular locus, but instead capturing variation of a genome-wide sample of markers. The more challenging scenario is the intermediate case in which a trait is controlled by both loci that are discernible from the polygenic background and the polygenic background itself (Rutkoski *et al*. 2014; Rice and Lipka 2019; DeWitt *et al*. 2021).

The ratio between the variance explained by the oligogenic and polygenic terms with the total genetic or phenotypic variance is likely a a major factor determining the cost-benefit of incorporating MAP, GP, or both into a breeding or diagnostic program. Modeling a individual loci can be advantageous when the proportion of the phenotypic and genetic variance explained by the locus is reasonably large and not partially captured by other markers in linkage disequilibrium (LD) with the target (Bernardo 2014; Rutkoski *et al*. 2014; Rice and Lipka 2019; Pincot *et al*. 2018, 2022). Ideally, the Feldmann et al 7 targeted markers should not fit the marker effect size distribution assumptions, e.g., that all marker effects contribute equally to the genomic variance and are drawn from the same distribution (Piepho 2009; Endelman 2011; Habier *et al*. 2007) and should not be in high LD with a large number of other markers. With ASV, researchers can accurately estimate these parameters directly in LMM analyses.

### Simulations confirm that ASV yields accurate estimates of all genetic variance components and ratios

As we show in our previous studies (Piepho 2019; Feldmann *et al*. 2021, 2022), ASV is ideal for estimating the variance explained by both single loci and GRMs. In our simulations, we included variation in population size, e.g., *n* = 500, 1, 000, and 1, 814, and replication of entries, e.g., *r* = 1, 2, and 4 for both outbred (Fig 1) and inbred populations (Fig 2). We can see that the same pattern that has emerged as in previous studies; the ASV approach yields accurate, unbiased estimates of variance components and variance component ratios from LMM analyses regardless of the constitution of the population or the study design. Even when there is only one replicate per entry (*r* = 1) all of the explanatory genetic terms are still accurately partitioned from the total variance. As *n* increased from 500 to 1, 814, the precision of estimates increased dramatically (the sampling variance decreases). Increasing *r* from 1 to 4 did not affect precision or accuracy of genomic and marker associated variances. However, increased numbers of replicates did improve the precision of residual variance components. This is because entries are replicated among plots (*n· r*), but markers and other genetic components are replicated among entries (*n*). Our simulations, in conjunction with our previous results (Piepho 2019; Feldmann *et al*. 2021, 2022), demonstrate that in most populations— human, animal, plant, or microbe—the average semivariance will yield accurate and easily interpreted estimates of different variance components.

**Figure 1.**
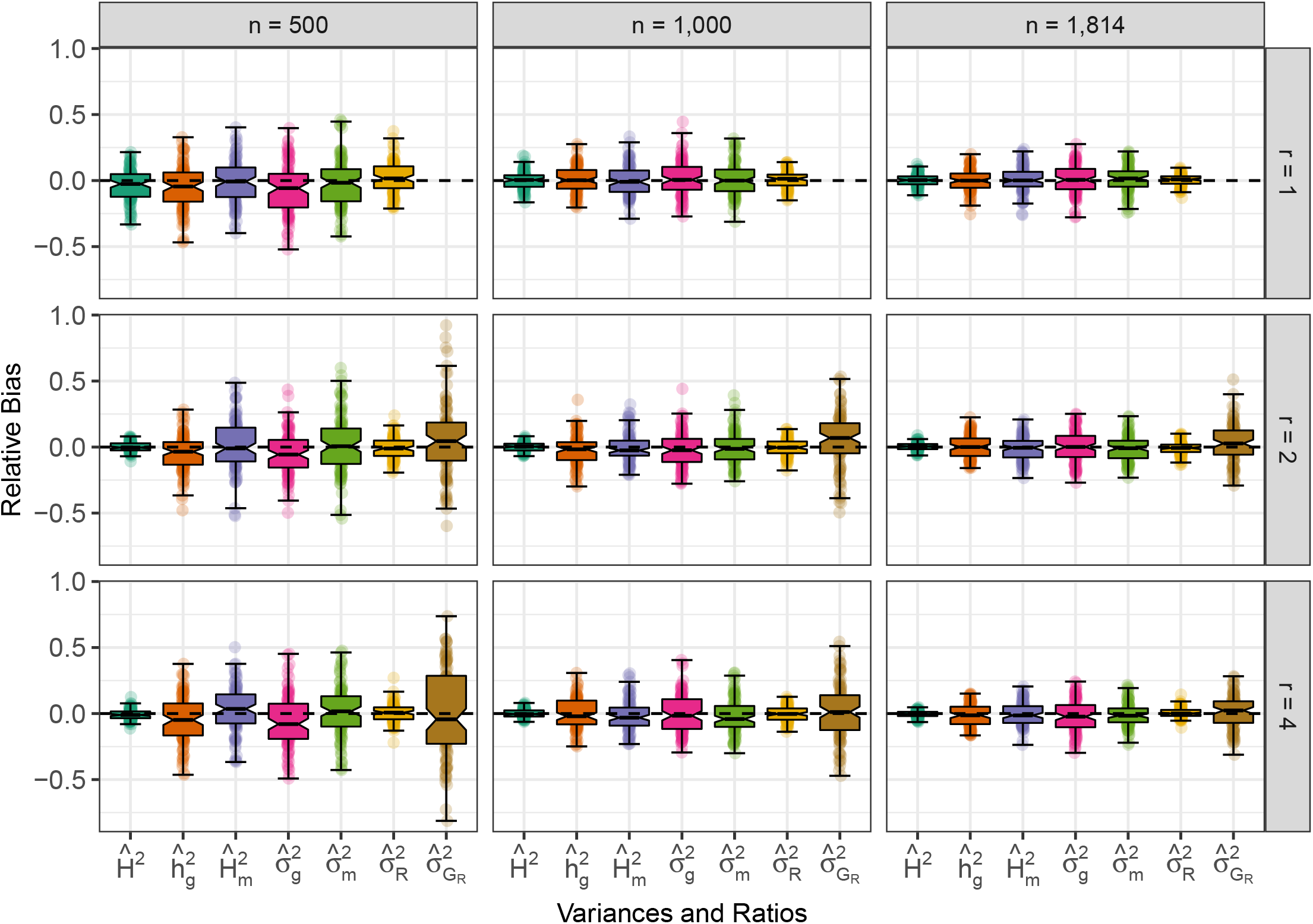
Effect of *n* and *r* on the relative bias of variance components and ratios in simulated outbred populations. Phenotypic observations were simulated for 100 samples with *n* = 500, 1, 000, and 1, 814 (left to right) genotyped for *m* = 5, 000 SNPs and the average heterozygosity *H* = 0.38. The relative bias of marker heritability, genomic heritability estimates 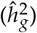, broad sense heritability, genomic variance, marker variance, residual genetic variance, and residual variance heritability when the number of replicates of each entry (*r*) = 1 (upper panel), 2 (middle panel), and 4 (lower panel). The upper and lower halves of each box correspond to the first and third quartiles (the 25th and 75th percentiles). The notch corresponds to the median (the 50th percentile). The upper whisker extends from the box to the highest value that is within 1.5 × *IQR* of the third quartile, where *IQR* is the inter-quartile range, or distance between the first and third quartiles. The lower whisker extends from the first quartile to the lowest value within 1.5 × *IQR* of the quartile. The dashed line in each plot is the true value from simulations.

**Figure 2.**
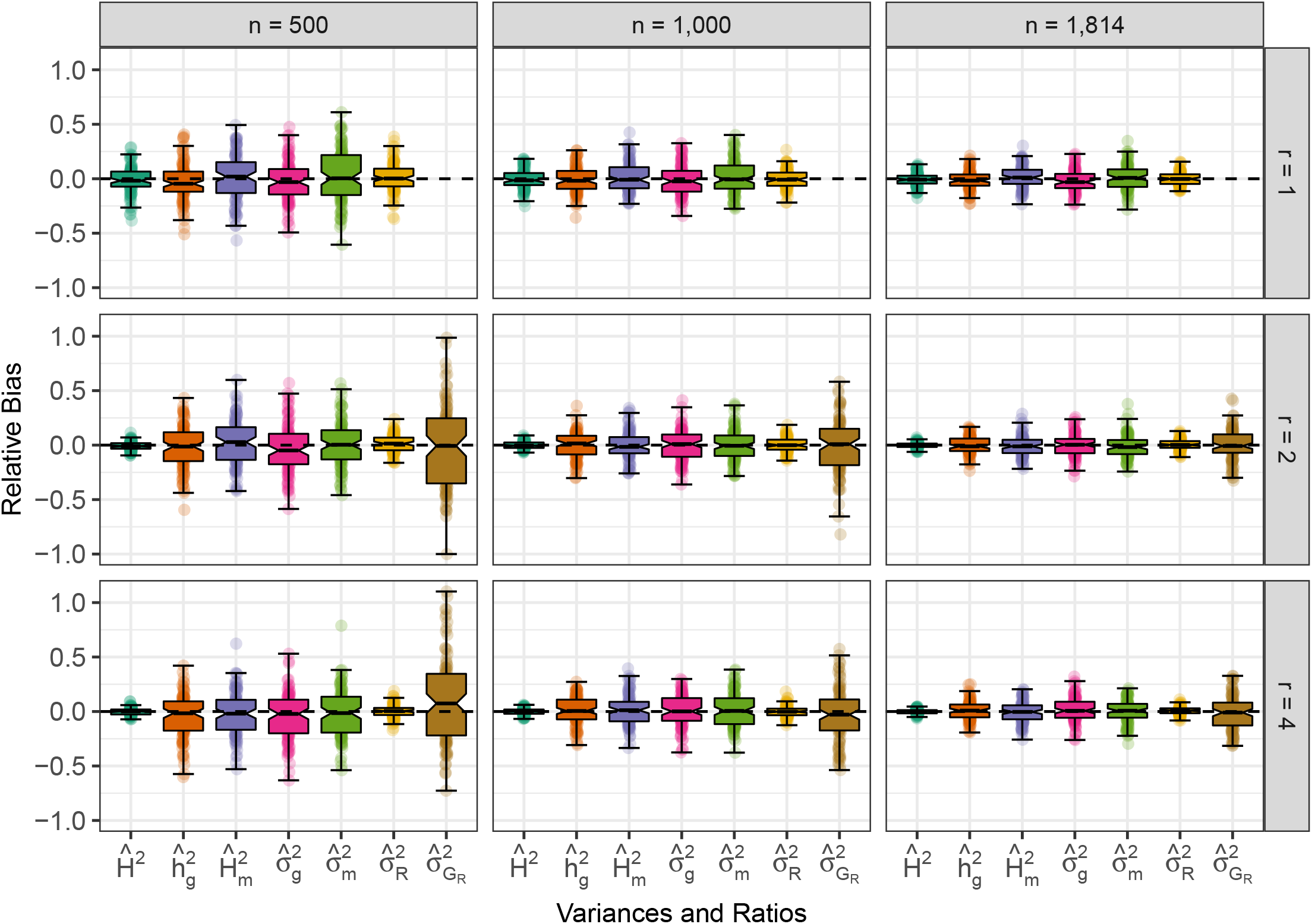
Effect of *n* and *r* on the relative bias of variance components and ratios in simulated inbred populations. Phenotypic observations were simulated for 100 samples with *n* = 500, 1, 000, and 1, 814 (left to right) genotyped for *m* = 5, 000 SNPs and the average heterozygosity *H* = 0. The relative bias of marker heritability, genomic heritability estimates 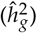, broad sense heritability, genomic variance, marker variance, residual genetic variance, and residual variance heritability when the number of replicates of each entry (*r*) = 1 (upper panel), 2 (middle panel), and 4 (lower panel). The upper and lower halves of each box correspond to the first and third quartiles (the 25th and 75th percentiles). The notch corresponds to the median (the 50th percentile). The upper whisker extends from the box to the highest value that is within 1.5 × *IQR* of the third quartile, where *IQR* is the inter-quartile range, or distance between the first and third quartiles. The lower whisker extends from the first quartile to the lowest value within 1.5 × *IQR* of the quartile. The dashed line in each plot is the true value from simulations.

**Figure 3.**
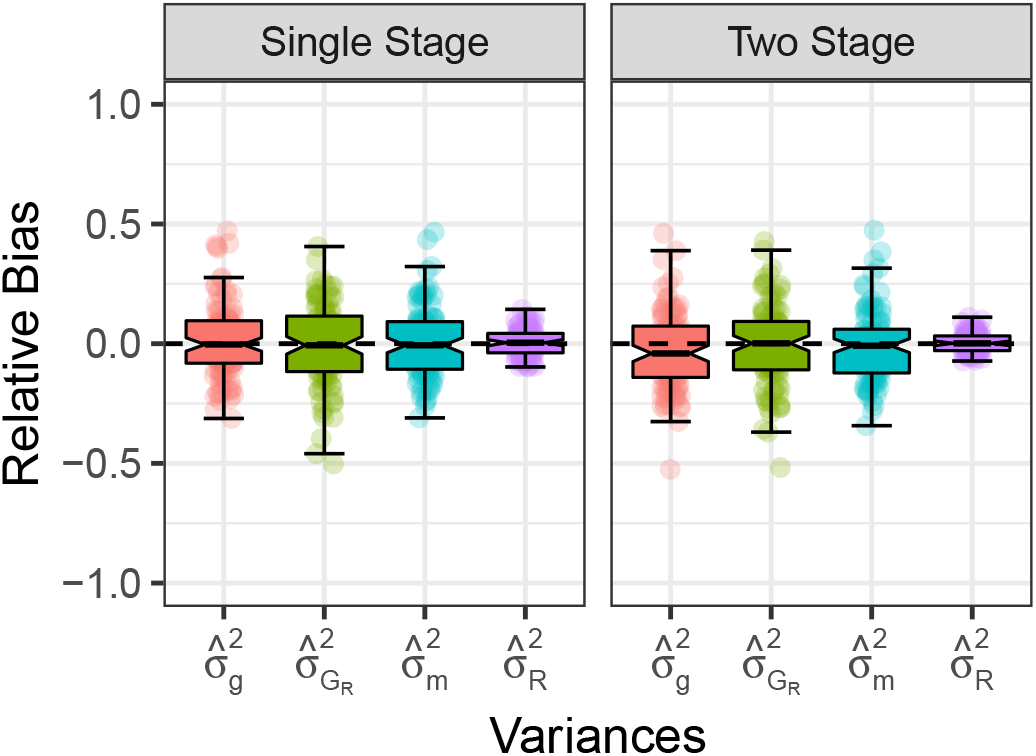
Single versus multi Stage analysis with two environments. The relative bias of genomic variance 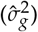, marker variance 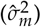, residual genetic variance 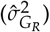, and residual variance 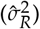analysed in a single stage (left panel) or in two stages (right panel). The upper and lower halves of each box correspond to the first and third quartiles (the 25th and 75th percentiles). The notch corresponds to the median (the 50th percentile). The upper whisker extends from the box to the highest value that is within 1.5 *IQR* of the third quartile, where *IQR* is the interquartile range, or distance between the first and third quartiles. The lower whisker extends from the first quartile to the lowest value within 1.5 *IQR* of the quartile. The dashed line in each plot is the true value from simulations.

### Average semivariance in quantitative genetics and beyond

ASV is a strategy that can be used for estimating and partitioning the total variance into components (Piepho 2019), such as the variance explained by loci and locus-locus (Feldmann *et al*. 2021) and the genomic variance (Feldmann *et al*. 2022). The approach we are suggesting shares some common threads with the current thinking in quantitative genetics, particularly as it relates to genomic relatedness, genomic heritability, and genomic prediction (VanRaden 2008; Yang *et al*. 2010; Kang *et al*. 2010; Habier *et al*. 2013; Hayes *et al*. 2009; Meuwissen *et al*. 2001; Isik *et al*. 2017; Zas and Sampedro 2015; Potti and Canal 2011; Roff and Fairbairn 2015; Swarts *et al*. 2021; Nietlisbach *et al*. 2016; Ulrich *et al*. 2021; Fan *et al*. 2021) but it also deviates from the classic quantitative genetic model conceptually in that it assumes that marker effects are random variables (Falconer and Mackay 1996; Lynch and Walsh 1998; Bernardo 2001). We have demonstrated that these are a statistically valid set of assumptions, even though they deviate from the classic quantitative genetics perspective.

ASV has several beneficial elements that make ASV a viable option for quantitative genetics, but more importantly, it is appropriate for any quantitative discipline where variance components are of interest from plant and microbial biology to psychology and infant research. Namely:

1. **The definitions of the variance components using average semivariance are additive and sum to the phenotypic variance**. This means that the LMM can be extended to incorporate all explanatory components, e.g., dominance, epistasis, transcriptomic, and will yield accurate VCEs for all terms (Nishio and Satoh 2014; Vitezica *et al*. 2017; Xiang *et al*. 2018; Krause *et al*. 2019). This is not necessarily true for all definitions of variance components (Piepho 2019).
2. **ASV is well suited for mutli-stage analyses** At the center of ASV, is the idea that the “entry mean” is the phenotype *per se*, and not the observations (Piepho 2019; Feldmann *et al*. 2022). One interpretation is that individuals, not observations, are the primary source of variation. ASV yields accurate estimates of the genetic and genomic variance components in unreplicated, or partially replicated, designs common in humans and agricultural plants and animals (Cullis *et al*. 2006; Moehring *et al*. 2014; Cullis *et al*. 2020; Butler *et al*. 2014; González-Barrios *et al*. 2019). ASV also yields accurate estimates in the twostage approaches to GP and GWAS in plants (Piepho *et al*. 2012; Damesa *et al*. 2017, 2019).
3. **ASV does not affect or impact the BLUPs or breeding value predictions**. ASV is only used to obtain accurate VCEs (Piepho 2019; Feldmann *et al*. 2022). It has been demonstrated that marker coding and different strategies for scaling and centering **Z** and **K** do not impact BLUPs or prediction accuracy (Strandén and Christensen 2011; Legarra 2016; Legarra *et al*. 2018), and, because ASV essentially works through a set of scalar coefficients determined by the experiment and population, this feature directly applies to this work.
4. **ASV works under many model assumptions in GLMM analyses** beyond the often-assumed variance-covariance structure in this study, e.g., 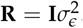. ASV can be applied to designs accounting for spatial structure through auto-regressive correlations or spline-models (Rodríguez-Álvarez *et al*. 2018; De Resende *et al*. 2006; Selle *et al*. 2019, 2020; Burgueño *et al*. 2000; Borges *et al*. 2019; Hoefler *et al*. 2020). ASV can also be applied to data sets where the observational units lead to non-normality of residuals; i.e., ordinal disease scores and proportion scores (Piepho 2019).

As substantiated by our simulations in this study and in the context of our previous work, ASV with REML estimation of the underlying variance components yields accurate estimates for oligo and polygenic effect, both individually and collectively, and BLUPs of the the additive and dominance effects of marker loci (Piepho 2019; Feldmann *et al*. 2021, 2022). ASV directly yields accurate estimates of genomic heritability in the observed population and can be used to adjust deviations that arise from other commonly used methods for calculating genomic relationships regardless of the population constitution, such as inbred lines and F_1_ hybrids, unstructured GWAS populations, or animal herds and flocks. We believe that **K**_*ASV*_ provides a powerful approach for directly estimating genomic heritability for the observed population regardless of study organism or experiment design (Visscher *et al*. 2006, 2007, 2008, 2010). In conclusion, our recommendation is that the average semi-variance approach be considered for general adoption by genetic researchers working in humans, microbes, or (un)domesticated plants and animals.

## Methods and Materials

### Computer Simulations

We generated 18 experiment designs with different population sizes of *n* = 500, 1, 000, and 1, 814 and number of clonal replicates per entry *r* = 1, 2, and 4 for outbred *H* = 0.38 and inbred *H* = 0.0 populations. Clonal replicates are a special case common in plant genetics of hybrid (e.g., maize, rice, and sorghum) cropping systems and in clonally propagated species (e.g., strawberry, potato, and apple). In all examples, 100 populations genotyped at *m* = 5, 000 loci. These 5, 000 SNPs were used to generate the purely additive polygenic background and one locus for the simple genetic effect. Marker genotypes, e.g., alleles, were drawn from a multivariate normal distribution with to replicate the population structure of the 1,814 mice from Valdar *et al*. (2006) using `MASS::mvrnorm()` and transformed such that the population was heterozygosity *H* = 0.38. We then estimated **K**_*ASV*_ and excluded the targeted locus from the calculation of **K**_*ASV*_. We also simulated residual genetic and residual effects each from a normal distribution with *µ* = 0 and 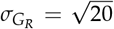and 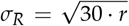using `stats::rnorm()`. A single explanatory locus was simulated with a segregation ratio of approximately 1 : 2 : 1 for AA:Aa:aa marker genotypes was simulated with *µ* = 0 and 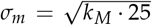 using `stats::rnorm()`. We did not control for the portion of additive vs dominance variance for the single marker. We simulated marker effects for all *m* = 5,000 loci following a normal distribution *µ* = 0 and 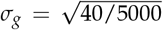. When multiplied by the centered marker genotypes and summed, the score is taken as the true additive genetic value *g* of each individual. For each simulated population we expressed LMM (1) using `asreml::asreml()` Butler (2021). In the second set of simulations, we used the same approach and same mean and variance parameters. However, in this example we simulated full inbred lines in the background polygenic markers (*H* = 0.0) and in the foreground markers, e.g., 1 : 0 : 1 for AA:Aa:aa. All plots are made with the `ggplot2` package Wickham (2016) in R 4.1.0 R Core Team (2020).

### Model statements in R/sommer v4.1.7

#### Incorporating One Target Locus into GBLUP

LMM (1) is expressed as:

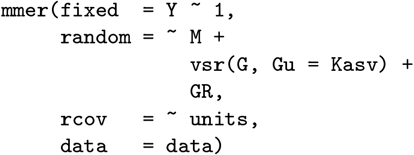

where `data` is a *n×* 4 matrix containing the phenotypic observations *Y*, a factor coding levels of *M*, a factor coding entries *G*, and a factor coding levels of *G*_*R*_. The variable `units` is inferred by `sommer::mmer()` and can be considered as a column with as many levels as rows in the data (Covarrubias-Pazaran 2016). The factor levels of *G* and *G*_*R*_ are equivalent.

The version of this model with *k*_*M*_ embedded is expressed as:

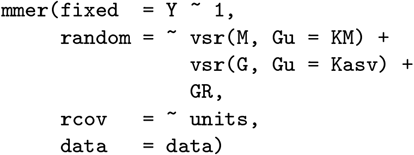

where `KM` is the matrix 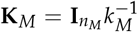. All other variables are the same as previously defined.

#### Incorporating Multiple Target Loci into GBLUP

LMM (8) is expressed as:

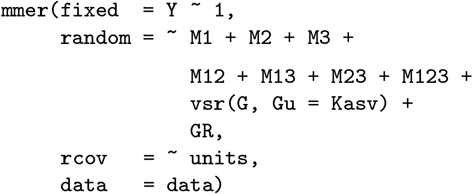

where `data` is a *n*× 10 matrix containing the phenotypic observations *Y*, seven columns corresponding to the marker effects and interactions, a factor coding entries *G*, and a factor coding levels of *G*_*R*_. The factor coding of *m*_*α*_ has three levels corresponding to *AA* : *Aa* : *aa* and a factor coding levels of *m*_*δ*_ has two levels corresponding to homozygous and heterozygous.

#### Partitioning Marker Variance into Additive and Dominance Components

LMM (9) is expressed as:

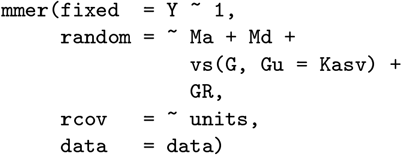

where `data` is a *n×* 5 matrix containing the phenotypic observations *Y*, a factor coding levels of *m*_*α*_, a factor coding levels of *m*_*δ*_, a factor coding entries *G*, and a factor coding levels of *G*_*R*_. The factor coding of *m*_*α*_ has three levels corresponding to *AA* : *Aa* : *aa* and a factor coding levels of *m*_*δ*_ has two levels corresponding to the genetic state—either homozygous or heterozygous.

#### Incorporating a Genomic Dominance Relationship Matrix into GBLUP

LMM (12) is expressed as:

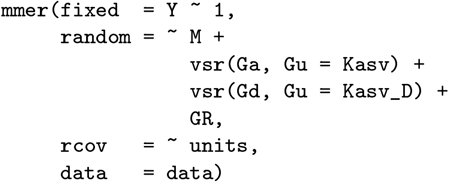

where `data` is a *n×* 5 matrix containing the phenotypic observations *Y*, a factor coding levels of *M*, and three factors coding entries, e.g., *G*_*α*_, *G*_*δ*_, and *G*_*R*_. The factor levels of *G*_*α*_, *G*_*δ*_, and *G*_*R*_ are equivalent.

#### Incorporating Stagewise Meta-analysis into GBLUP

LMM (14) is expressed as:

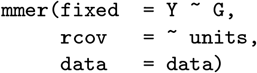

where `data` is a *n×* 2 matrix containing the phenotypic observations *Y* and one factor coding *G* for the entry ID. Blocks and other within location design elements can be incorporated as random effects using the `random =` syntax. In sommer, **R**_*e*_s are obtained from each location as the ‘VarBeta’ matrix in the `sommer::mmer()` output. Specially, ‘VarBeta’ is the name of the model estimated variance covariance matrix among entry means in sommer. The **R**_*e*_s are then bound corner-to-corner, which is accomplished using `sommer::adiag1()` to obtain **Ω**. We then take the inverse of **Ω** using `base::solve()`.

The LMM for stage 2 (16) is expressed as:

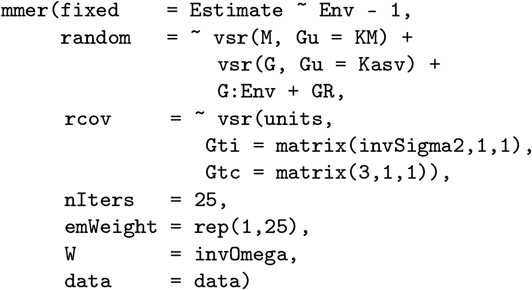

where where `data` is a *n×* 5 matrix containing the adjusted entry means from stage 1 *Y*, a factor coding levels of *M*, two equivalent factors coding entries, e.g., *G* and *G*_*R*_, and one factor coding environments *Env*. In this approach, we must fix the residual variance component equal to 1 so that the residual so that all the scaling of the `invOmega` = **Ω**^*−*1^ is unaffected by the model estimation process. Within the `vs()` argument, the `Gti()` and `Gtc()` arguments are used to set the initial value of the variance component equal to the inverse of the variance among adjusted entry means 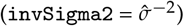 and to constrain the variance component estimation to a fixed value by setting the first argument equal to 3 (Covarrubias-Pazaran 2022). In this example we use 25 iterations of 100% expectation-maximization algorithm; however, the EM and NR methods can be exchanged or averaged, by changing the `emWeight` argument.

## Data Availability

Zenodo repository coming soon. For now, code is available by request.

## Author Contributions

**Conceptualization:** MJF, HPP **Data curation:** MJF **Formal Analysis:** MJF **Funding Acquisition:** HPP **Investigation:** MJF, HPP **Methodology:** MJF, GCP **Project administration:** MJF, HPP **Resources:** HPP **Software:** MJF, GCP **Supervision:** MJF, HPP **Validation:** MJF **Visualization:** MJF **Writing – original draft preparation:** MJF, HPP **Writing – review & editing:** MJF, HPP, GCP

